# Neuronally differentiated endothelial cell subtype regulates organ blood flow and immune balance

**DOI:** 10.1101/2024.09.30.615824

**Authors:** Georgina Gyarmati, Ruslan Rust, Alejandra Becerra Calderon, Audrey Izuhara, Greta Trogen, Sachin Deepak, Yibu Chen, Seth Walter Ruffins, Jason A. Junge, Berislav V. Zlokovic, Scott Fraser, János Peti-Peterdi

**Author notes:** Corresponding authors: Georgina Gyarmati, MD, PhD, MPH, Janos Peti-Peterdi, MD, PhD, Department of Physiology and Neuroscience, Zilkha Neurogenetic Institute, University of Southern California, 1501 San Pablo Street, Room ZNI 335, Los Angeles, California 90033.

## Abstract

Vascular endothelial cells (ECs) perform key pleiotropic functions to maintain body homeostasis via the regulation of organ blood flow, vascular permeability, tissue growth and inflammation, and angiogenesis. Recent transcriptomic studies uncovered many EC subtypes across organs; however their specific functions are incompletely understood. Here we identified and characterized a novel, minority subtype of scattered ECs with a well-defined arteriovenous zonal localization exclusively in small resistance (strain) arterioles, and with the highest density in the brain>retina>kidney. Due to their expression of both endothelial and neuron-like functional and gene transcriptomic signatures, they were termed neuro-endothelial cells (NECs). High resolution single-cell transcriptome analysis of mouse brain and kidney ECs identified neuronal nitric oxide synthase (Nos1) and cytokine-like 1 (Cytl1) as top NEC biomarkers. Intravital multiphoton imaging of optogenetic mouse models with NEC gain/loss-of-function revealed NEC and Nos1-dependent vasodilation/vasoconstriction of intact brain and kidney arterioles and elevation/reduction in blood flow. Silencing NEC Nos1 and Cytl1 gene expression in vivo caused marked segmental arteriolar vasoconstrictions, reductions in vascular density and organ blood flow, increased vascular permeability and immune cell homing. Cytl1 administration triggered vasodilation and increased blood flow acutely, and increased capillary density and clonal EC remodeling chronically. NECs play major vasodilatory, angiogenic and anti-inflammatory functions that may be therapeutically targeted for vascular and inflammatory diseases.

## Introduction

The endothelium plays a pivotal role in vascular function and vascular homeostasis, including regulation of vascular tone, permeability, inflammation, and angiogenesis. Whereas vascular endothelial cells (ECs) are traditionally considered a single cell type, they exhibit considerable structural, phenotypic, and functional heterogeneity depending on the tissue in which they reside^1–3^. In the past few years, numerous studies performed high-resolution transcriptome analysis of ECs in different cell states and established remarkable EC heterogeneity in distinct tissues from adult mice and humans^1,4,5^. These works identified substantial organotypic differences between brain, lung, heart, kidney, liver, and other organs’ vascular cell types and established EC heterogeneity features based on arteriovenous zonation. In addition, these studies provided powerful discovery tools in the form of publicly available and online searchable single-cell transcriptome atlases, most commonly of the brain vasculature in normal physiological or disease conditions^5–8^. Based on these new scRNA seq transcriptome data, there is general agreement that ECs display a high level of heterogeneity that correspond to distinct vascular beds in terms of organ/tissue type, arterial-capillary-venous zonation (e.g. Vegfc for arterial ECs)^5^, metabolic states (e.g. Txnip for arterial ECs)^6^, and sex differences (e.g. Lars2 for male ECs)^4^. In addition, recent work established the high-precision and genome-scale gene expression landscape of early embryonic vascular ECs and confirmed the presence of significant endothelial heterogeneity already during early vascular development^9^.

In addition to their physiological roles in maintaining vascular health and tissue homeostasis, EC dysfunction is a hallmark of vascular diseases^10–12^. Tissue-specific EC dysfunction is known to play important roles in the development of reduced blood flow and a spectrum of organ diseases. In the blood–brain barrier (BBB), for example, ECs are bound by tight junctions to maintain a highly selective, low-permeability barrier. BBB endothelial dysfunction can lead to Alzheimer disease, epilepsy, and multiple sclerosis^13–15^. Glomerular ECs in the kidney are critically important component of the glomerular filtration barrier and restrict albumin filtration, and albuminuria is strongly associated with EC dysfunction^16^. The cardiac endothelium plays a crucial role in promoting cardiomyocyte proliferation and maturation via paracrine signaling^17,18^, while the secretion of angiocrine factors from pulmonary and liver sinusoidal ECs is critical in modulating lung^19^ and hepatic regeneration^20^. Thus, understanding tissue-specific EC functionality is critical for treating a wide range of human diseases.

The regulation of blood flow distribution across different organs’ vascular beds, and vascular autoregulation are critically important physiological control mechanisms that maintain tissue perfusion in response to stress, including varying metabolic needs, hypovolemia and blood pressure alterations. These vascular regulatory mechanisms primarily involve complex adjustments of the tone of resistance (strain) arterioles in the peripheral microcirculation. Nitric oxide (NO) produced by ECs has been generally considered as the primary vasodilator and a regulator of vascular functions. Although classically, endothelial NO synthase (eNOS, Nos3) is considered the main NO synthase isoform involved in vascular regulation, a body of work from animal as well as human studies has suggested that neuronal NO synthase (nNOS, Nos1) is also expressed in the vascular endothelium, and Nos1-derived NO contributes to the maintenance of cardiovascular homeostasis^21–23^. However, the cellular source of Nos1-derived NO and the detailed molecular mechanisms have been elusive.

The present study aimed to improve our understanding of organ-specific EC heterogeneity at the cell and molecular level, and to identify and characterize the Nos1^+^ EC lineage including their function in the healthy brain and kidney and relevant to vascular disease.

## Results

### Genetic cell fate tracking of the Nos1 cell lineage

Recent work with tamoxifen-inducible Nos1-CreERT2 mice and floxed mTmG reporter (Nos1-mTmG mice) for genetic labeling of the Nos1^+^ cell lineage in the kidney identified the neuronally differentiated renal cell type of the macula densa^24^. However, prolonged treatment with maximum dose tamoxifen uncovered a few additional Nos1^+^ cells in the renal cortex in close proximity to glomeruli. Systematic genetic lineage tracing of Nos1^+^ cells in all organs using this mouse model identified a novel, minority subtype of scattered Nos1^+^ ECs with a well-defined arteriovenous zonal localization exclusively in small resistance (strain) arterioles, and none in capillaries or veins (Fig. 1 and Supplemental Fig. 1). Due to the expression of both endothelial (e.g. CD31) and neuron-like (e.g. Nos1) structural, functional and gene transcriptomic signatures (Figs. 1-2, and Supplemental Figs. 1-2), these Nos1^+^ endothelial cells were termed neuro-endothelial cells (NECs). Based on histological analysis (Fig. 1A, and Supplemental Fig. 1C) and subsequent confirmation by flow cytometry (Fig. 1B and Supplemental Fig. 1B), NECs were found with the highest density in the brain>retina>kidney (1-4% of all ECs)(Supplemental Movie 1), and with lower but detectable density in the heart and pancreas (0.1-0.3%). Compared to other ECs, NECs featured a more elongated morphology with long, upstream cell processes (pointing against the direction of blood flow), and numerous small membrane protrusions (Fig. 1C). Additional genetic cell fate tracking of the Nos1 lineage, either over extended periods of time (6 months) after a single tamoxifen induction given at weaning, or after continuous tamoxifen induction for two months combined with Nos1 immunolabeling, confirmed the same scattered, low-density NEC distribution in resistance arterioles, and the exclusively NEC-specific Nos1 protein expression (Supplemental Fig. 1A). These studies confirmed that NECs represent a permanent cell type with terminal neuronal differentiation rather than a transient cell state. Importantly, no other vascular cell types were labeled in long-term fate tracking studies including no other ECs in arterioles, capillaries and venules, and vascular smooth muscle cells (VSMC) (Supplemental Fig. 1A), suggesting that NECs do not function as vascular precursor cells. In addition to mice, Nos1^+^ NECs were found in human brain and kidney arterioles (Supplemental Fig. 1C).

**Figure 1.**
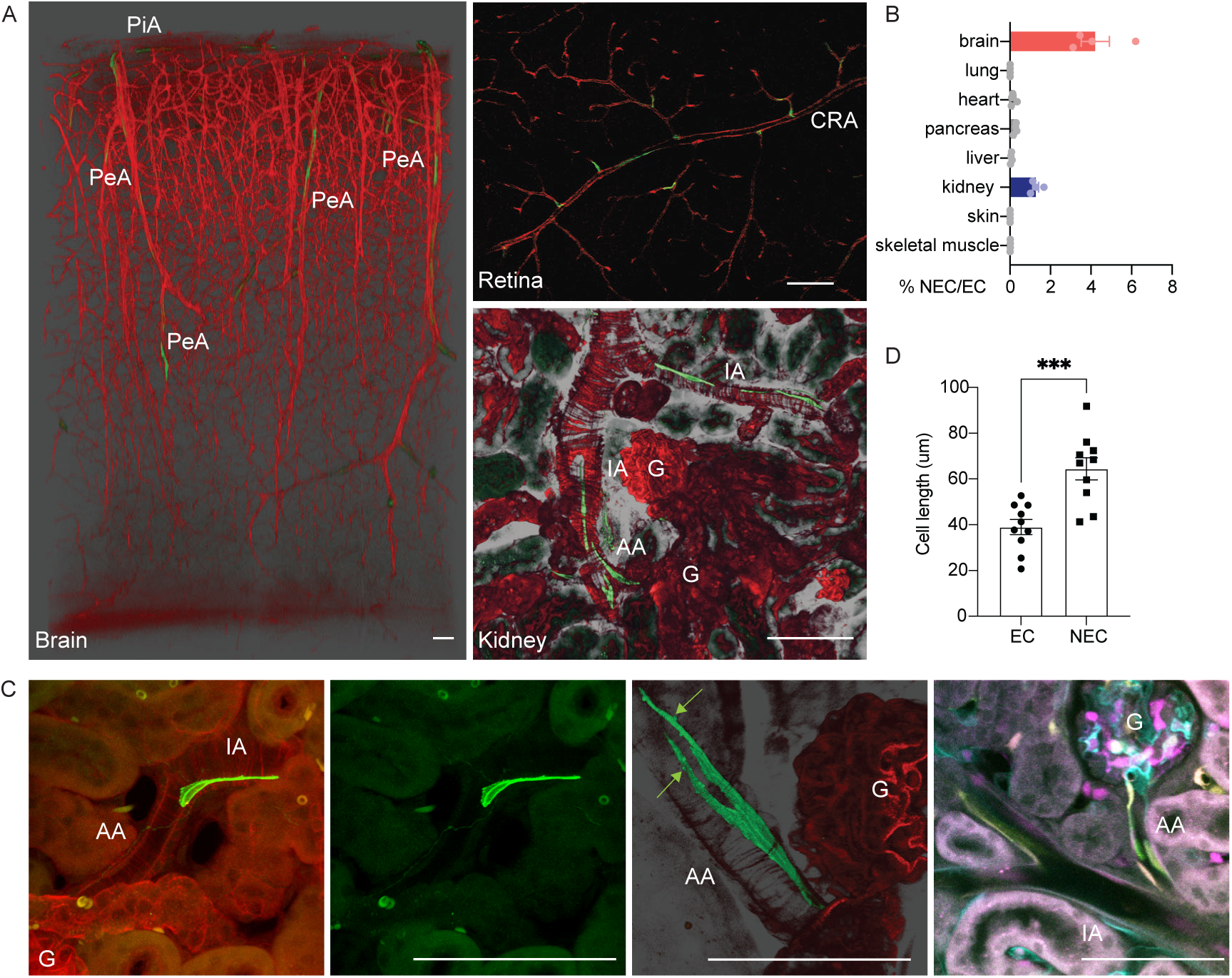
NEC identification, localization, and features in the brain, retina, and kidney vasculature. **(A)** Representative 3D and 2D light-sheet or multiphoton fluorescence images from fixed, optically cleared whole-mount Nos1-mTmG mouse brain, retina, and kidney. Note the presence of a few scattered NECs (green) in pial (PiA) and penetrating arterioles (PeA) in the brain, in inner retina arteriole branches supplied by the central retinal artery (CRA), and in interlobular (IA) and afferent arterioles (AA) in the kidney. All other cell types are labeled by the genetic reporter tdTomato (red). Arterioles are identified by the typical ring-like and intensely tdTomato^/^ vascular smooth muscle cells. G: glomerulus. **(B)** NEC density (normalized to the number of all ECs) across organs quantified by flow cytometry or histology (n=4). **(C)** Cell morphology features of NECs vs ECs on 3D or 2D images of Nos1-mTmG (3 left panels) or Cdh5-Confetti (right) mouse kidney. Note the more elongated NECs with long, upstream cell processes and numerous small membrane protrusions (arrows). Bars are 100 µm. **(D)** Summary of NEC vs EC cell length (n=10 cells from n=4 kidneys each). Data are mean ± SEM, ***p<0.001 with t-test.

**Figure 2.**
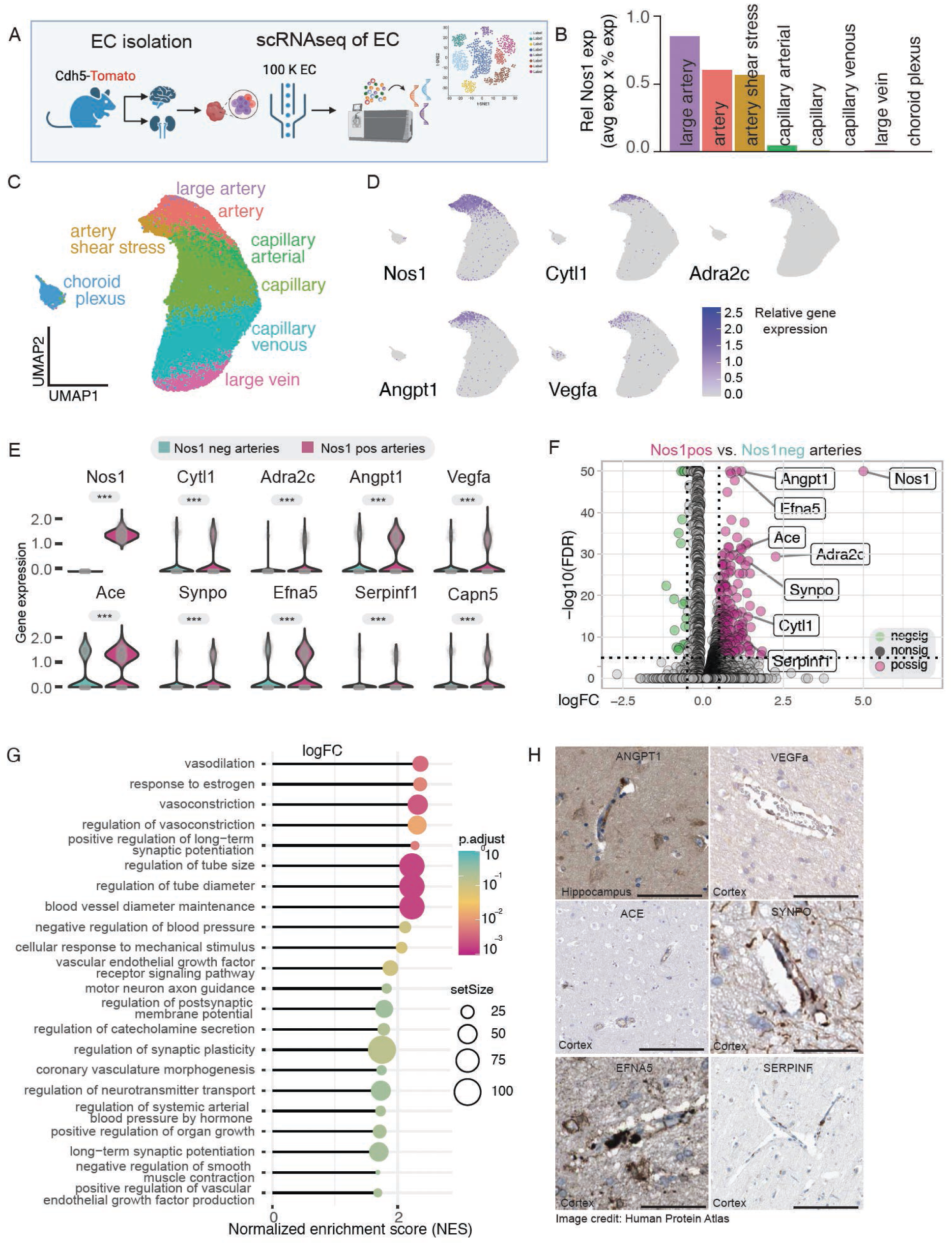
Transcriptome analysis of brain NECs. **(A)** Illustration of the workflow. **(B)** Relative expression of Nos1 in the main vascular segments’ EC subtypes according to arterio-venous zonation. **(C-D)** UMAP visualization of scRNA seq-based transcriptome analysis of about 100K mouse brain ECs with color-coded arterio-venous zonation (C) and the expression of top NEC-specific DEGs including *Nos1* and *Cytl1* (D). Note the similar pattern of high *Nos1, Cytl1, Adra2c, Angpt1*, and *Vegfa* expression only in arteriolar ECs (purple). **(E-F)** Violin (E) and volcano (F) plot of the expression of top NEC-specific DEGs in Nos1^+^ vs Nos1^-^ arteriolar ECs. **(G)** Gene set enrichment analysis with the top, most significant NEC biological functions in the brain. **(H)** Immunohistochemistry validation of the expression of top enriched mouse NEC-specific genes in the human brain. Data are from the Human Protein Atlas (HPA); images available from https://www.proteinatlas.org/ENSG00000089250-NOS1/tissue/cerebral+cortex#img Scale bar is 100 μm.

### NEC transcriptome analysis

As the initial approach to characterize NECs and to understand their molecular signature, high-resolution (100K reads/cell) single-cell RNA sequencing and transcriptome analysis was performed from 100K isolated ECs each from Cdh5-tdTomato mouse brain and kidney (Figs. 2-3). EC subtypes showing arterial, capillary and venous characteristics were identified based on well-known, published markers in both organs (Supplemental Fig. 2). Accordingly, graph-based clustering and UMAP visualization confirmed several different arterio-venous clusters of ECs in both the brain and kidney (Figs. 2B,C-3A,B). Importantly, this approach identified Nos1^+^ NECs as a distinct cell cluster within arterial ECs (Figs. 2B,D-3A,C) confirming the image-based cell fate tracking data (Fig. 1). NECs localized only to pre-glomerular arterioles in the kidney (Fig. 3A). Differentially expressed genes (DEGs), biomarker and gene set enrichment analysis of Nos1^+^ vs Nos1^-^ arteriolar ECs identified several NEC-specific vasoactivity (*Nos1, Adra2c, Ace*), and angiogenesis regulating (*Vegfa, Cytl1, Angpt1, Serpinf1*), chemotactic and immune regulatory (*Cytl1, F2rl1, C1ra*), tissue growth, extracellular matrix (ECM) remodeling and patterning factors (*Lox, Bmp2*) suggesting the role of NECs in regulating distal organ blood flow, vascular density, immune balance and inflammation, and vascular remodeling (Figs. 2E-G, 3D-I, Supplemental

**Figure 3.**
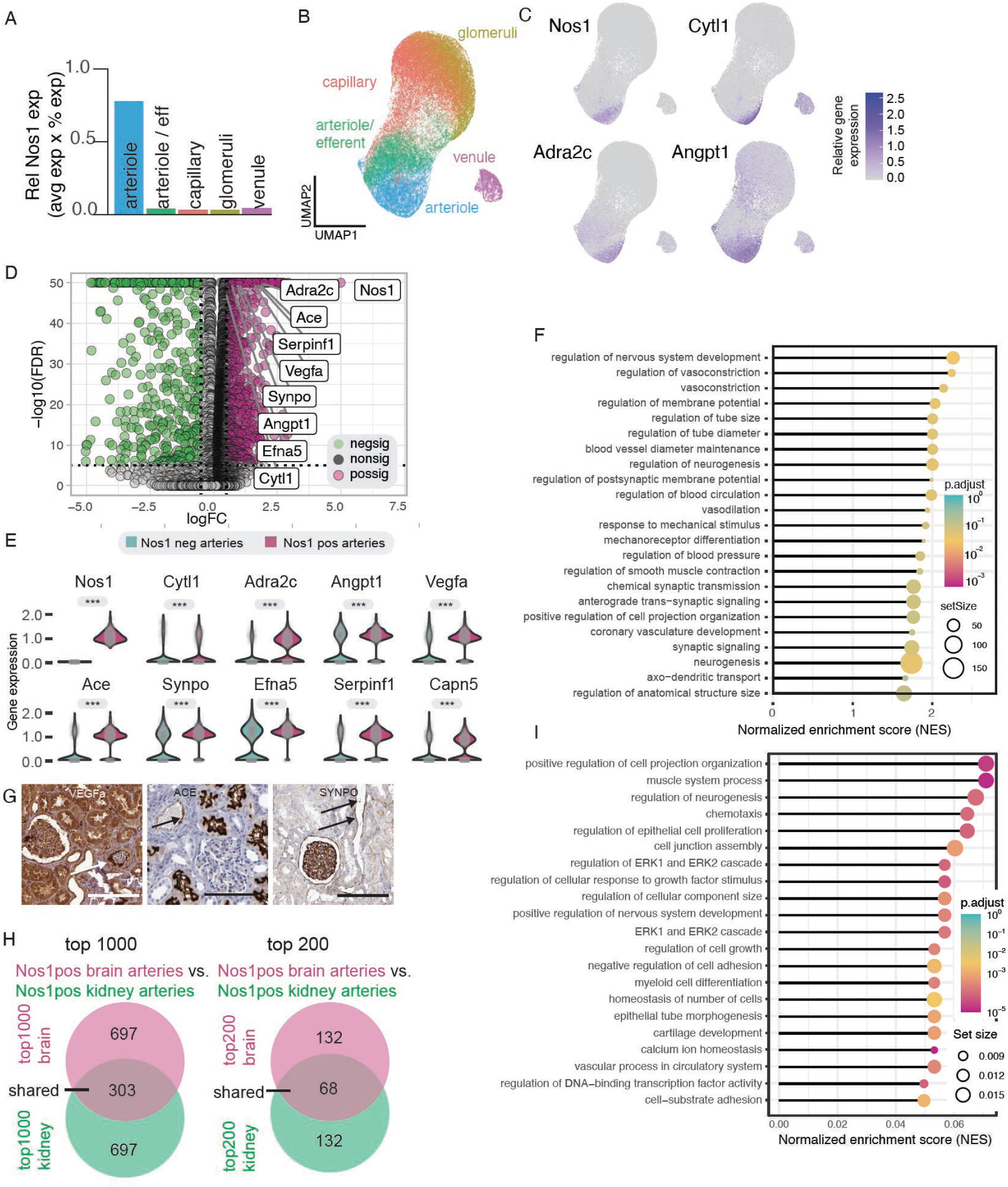
Transcriptome analysis of kidney NECs. **(A)** Relative expression of Nos1 in the main vascular segments’ EC subtypes according to arterio-venous zonation. **(B-C)** UMAP visualization of scRNA seq-based transcriptome analysis of about 100K mouse kidney ECs with color-coded arterio-venous zonation (B) and the expression of top NEC-specific DEGs including *Nos1* and *Cytl1* (C). Note the similar pattern of high *Nos1, Cytl1, Adra2c*, and *Angpt1* expression only in arteriolar ECs (purple). **(D-E)** Volcano (D) and violin (E) plot of the expression of top NEC-specific DEGs in Nos1^+^ vs Nos1^-^ arteriolar ECs. **(F)** Gene set enrichment analysis with the top, most significant NEC biological functions in the kidney. **(G)** Immunohistochemistry validation of the expression of top enriched mouse NEC-specific genes in the human kidney. Data are from the Human Protein Atlas (HPA); images available from https://www.proteinatlas.org/ENSG00000089250-NOS1/tissue/kidney#img Scale bar is 100 μm. **(H)** Homology of top NEC-specific DEGs identified from the brain and kidney. **(I)** Gene set enrichment analysis with the top shared genes in Nos1^+^ arteries in brain and kidney.

Table 1). Additional top NEC-specific biological functions included response to estrogen, mechanical stimuli (mechanosensation), and several neuron-like functions such as synaptic signaling and regulation, axon guidance, cell projection organization, and neurogenesis (Figs. 2G and 3F, I). The expression of some of the newly identified NEC-specific genes was validated and translated to the human brain and kidney on the protein level (Figs. 2H, 3G). The top NEC-specific DEGs identified from the brain and kidney showed high level of homology (Fig. 3H-I).

### Optogenetic modulation of NEC function

Following up on the transcriptome data, we next aimed to functionally test whether NECs regulate the vasoactivity of brain and kidney resistance arterioles. NEC-specific optogenetic gain-of-function (NEC-Ai27 mice with Nos1-driven expression of the improved channelrhodopsin-2/tdTomato fusion protein hChR2(H134R)/tdT in NECs) and loss-of-function (NEC-Ai39 mice with Nos1-driven expression of the improved halorhodopsin/EYFP fusion protein eNpHR3.0/EYFP) mouse models were established to measure alterations in arteriolar hemodynamics and organ blood flow in response to NEC optogenetic modulation (Fig. 4A). After validation of the NEC-Ai27 and 39 mouse models (Supplemental Fig. 3), quantitative imaging of resistance arterioles was performed in the intact brain using intravital multiphoton microcopy (MPM) and blue/red light stimulation. Using NEC-specific region of interest (ROI) scanning, single NEC stimulation (blue light in NEC-Ai27 mice) and inhibition (red light in NEC-Ai39 mice) revealed substantial increases and reductions in vessel diameter and organ blood flow, respectively (Fig. 4B-D) (Supplemental Movie 2). Pre-treatment with the Nos1 inhibitor 7-NI blocked the blue light-induced NEC NO synthesis and vasodilatations (Fig. 4C), confirming the regulatory role of NEC Nos1-derived NO in vascular contractility and organ blood flow.

**Figure 4.**
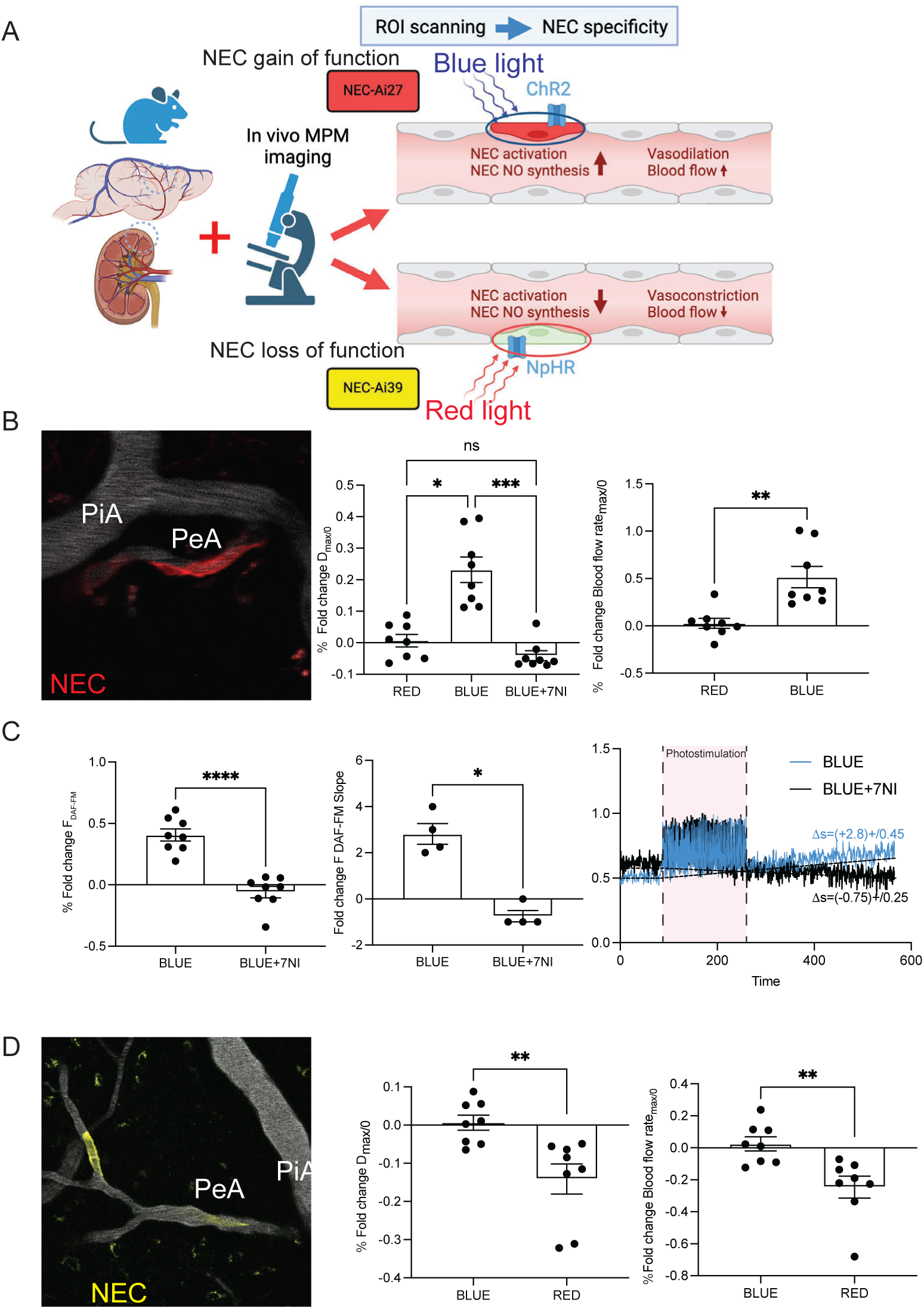
Intravital MPM imaging of the hemodynamic effects of acute single NEC optogenetic stimulation and inhibition. **(A)** Illustration of the optogenetic study design. **(B)** Representative intravital MPM image of a single NEC in a NEC-Ai27 mouse brain resistance arteriole (red due to Ai27 tdTomato reporter, left). Statistical summary of the relative changes in vessel diameter (center) and blood flow (right) normalized to baseline in response to NEC stimulation by blue light (n=8). **(C)** Representative time-lapse recordings (left) and statistical summary of NEC relative DAF-FM fluorescence intensity (center) and slope (right) at baseline, during, and after NEC photostimulation by blue light with (black line) or without (blue line) pretreatment with the Nos1 selective inhibitor 7NI (20mg/kg iv)(n=8). **(D)** Representative intravital MPM image of a single NEC in a NEC-Ai39 mouse brain resistance arteriole (yellow due to Ai39 EYFP reporter, left). Statistical summary of the relative changes in vessel diameter (center) and blood flow (right) normalized to baseline in response to NEC inhibition by red light (n=8). PiA: pial arteries, PeA: penetrating arterioles. Scale bars are 100 μm. Data are mean ± SEM, ns: not significant, *,**,***,****P<0.05, 0.01, 001, 0.001, t-test and ANOVA followed by Tukey’s test.

MPM imaging of intact brain and kidney arterioles in Cdh5-GCaMP6f/tdTomato mice revealed marked and direct NEC-specific calcium responses to acute norepinephrine challenge (Supplemental Fig. 3B) validating the functional expression of the Adra2c receptor in NECs (Figs. 2D-F, and 3C-E).

### Silencing NEC Nos1 and Cytl1 gene expression

To specifically test the functional role of NEC Nos1 and Cytl1 in brain and kidney arterioles in vivo, we next performed silencing of NEC Nos1 and Cytl1 gene expression in vivo using iv injections of 2X10^11^ GC/kg endothelial targeting AAV2-QUADYF serotype containing either control GFP RNA (NEC-WT), Nos1 shRNA (NEC-Nos1 knockdown (KD)), or Cytl1 shRNA (NEC-Cytl1 KD) (Fig. 5A). Histological analysis of endothelium-specific GFP labeling and Nos1 and Cytl1 RNAscope confirmed and validated the EC/NEC-specific targeting approach and highly efficient NEC Nos1 (76%) and Cytl1 (88%) gene silencing (Supplemental Fig. 4). General phenotyping revealed increased blood pressure in NEC-Nos1KD mice (Supplemental Fig. 4). Intravital MPM imaging of the brain and kidney 1 week after AAV2 injections found multiple severe and segmental, sphincter-like vasoconstrictions along all resistance vessels, and marked reductions in blood flow in both NEC-Nos1 KD and NEC-Cytl1 KD mice compared to NEC-WT (Fig. 5B). In addition, NEC-Cytl1 KD mice featured increased vascular permeability to plasma albumin compared to NEC-WT (Fig. 5C), and capillary density and vessel diameters were reduced in both NEC-Nos1 KD and NEC-Cytl1 KD compared to NEC-WT (Fig. 5D). In addition to these hemodynamic changes, MPM imaging of the plasma marker observed numerous negatively labeled cells sticking and rolling in the vascular lumen in NEC-Nos1 and Cytl1 KD animals. Specific labeling of all immune cells found high density of homing CD45^+^ immune cells in the arteriolar/capillary/venule lumen in NEC-Nos1KD and NEC-Cytl1KD mice in contrast to NEC-WT (Fig. 5E).

**Figure 5.**
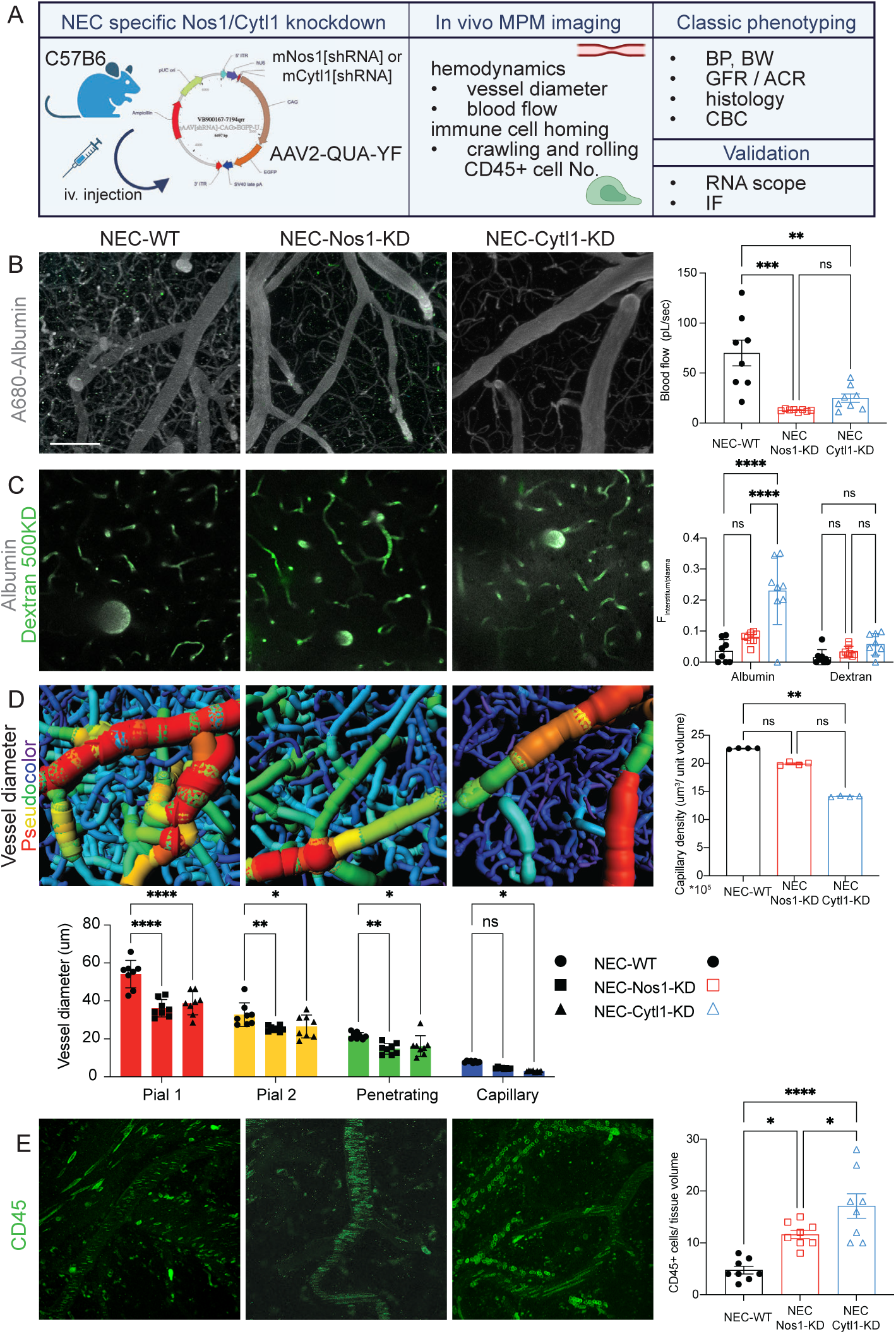
Vascular and immune cell homing effects of silencing NEC Nos1 and Cytl1 gene expression in vivo. **(A)** Illustration of the workflow and study design of NEC-specific Nos1 and Cytl1 knockdown and subsequent phenotyping. **(B-E)** Representative in vivo MPM images of brain resistance arterioles in control NEC-WT (AAV2-QUADYF-GFP RNA injected), NEC-Nos1 knockdown (KD) (AAV2-QUADYF-Nos1 shRNA injected), and NEC-Cytl1KD mice (AAV2-QUADYF-Cytl1 shRNA injected) mice, and statistical summary (right and bottom) of arteriolar blood flow (B), iv injected albumin-Alexa Fluor 680 and 500 kDa dextran-Alexa Fluor 488 permeability (C), capillary density and vessel diameter (D, image processing and filament tracing by Imaris), and the density of homing CD45^+^ immune cells labeled by iv injected anti-CD45-Alexa Fluor 488 antibodies (E). Note the multiple segmental sphincter-like vasoconstrictions (B, arrows), high interstitial albumin content along the resistance arterioles (C, arrows), and the high density of homing CD45^+^ cells in the arteriolar/capillary/venule lumen (E) in NEC-Nos1KD and NEC-Cytl1KD mice in contrast to control NEC-WT. Scale bar is 100 μm. Data are mean ± SEM, ns: not significant, *,**,***,****P<0.05, 0.01, 001, 0.001, ANOVA followed by Tukey’s test.

### Acute/chronic treatment with recombinant Cytl1

To test the therapeutic potential of mouse recombinant Cytl1 administration, intravital MPM imaging of brain and kidney resistance arterioles was performed in control healthy conditions and in the UUO model of kidney fibrosis and endothelial dysfunction^25^. Acute treatment of Cdh5-GT mice with Cytl1 (20µg/kg iv) resulted in marked vasodilatation and increased blood flow, and elevations in EC calcium in brain and kidney resistance arterioles (Fig. 6A-B, and Supplemental Movie 3). These vasculoprotective effects were preserved in disease conditions (Fig. 6B). In addition, chronic treatment with daily Cytl1 injections (20µg/kg BW/day sc. for 5 consecutive days) in Cdh5-Confetti mice induced angiogenesis and clonal endothelial remodeling evidenced by the increased number of Confetti^+^ ECs and unicolor clonal progenies in brain and kidney tissues (Fig. 6C).

**Figure 6.**
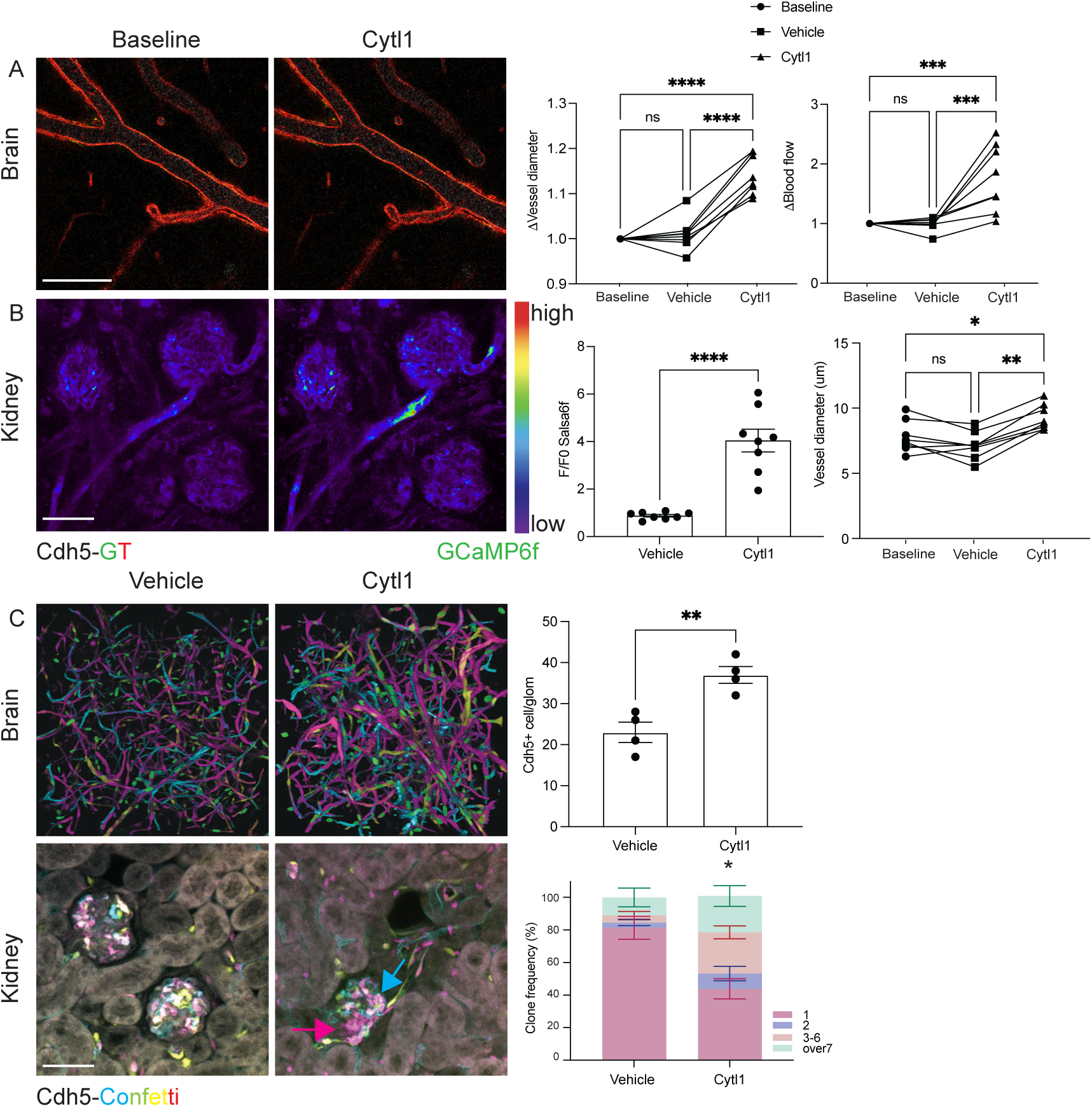
Acute and chronic effects of recombinant Cytl1 treatment in vivo. (A-B) Intravital MPM images of brain and kidney resistance arterioles in control healthy (A) and in UUO model of kidney fibrosis (B) Cdh5-GCaMP6f/tdTomato (GT) mice at baseline before (left) and within 3 minutes of acute iv injections of 20µg/kg BW mouse recombinant Cytl1 (center). Summary of Cytl1-induced changes in arteriole diameter and blood flow (A) and EC calcium (B)(right)(n=6 each). Note the dilated vessel lumen (A) and increased EC calcium in both afferent (AA)/efferent arterioles (EA) and glomerular (G) capillaries (B) in response to Cytl1. **(C)** Histological images of whole-mount cleared brain and kidney tissues of Cdh5-Confetti mice with EC genetic cell fate tracking and clonal remodeling based on multicolor Confetti^/^ cell distribution in control (left) and after chronic Cytl1 treatment for 1 week (20µg/kg BW/day sc. for 5 consecutive days)(center). Note the presence of unicolor cell clones (arrows) after Cytl1 treatment in contrast to the stochastic multicolor single-cell distribution in control. Angiogenesis and clonality analysis of tissue remodeling by the number of Confetti^/^ cells/glomerulus (top) and changes in the relative frequency of 1, 2, 3-6, and over 7 unicolor Confetti^/^ cells/clone (bottom)(n=4 each). Scale bars are 100 µm. Data are mean ± SEM, *,**,***,****P<0.05, 0.01, 001, 0.001, t-test.

**Figure 7.**
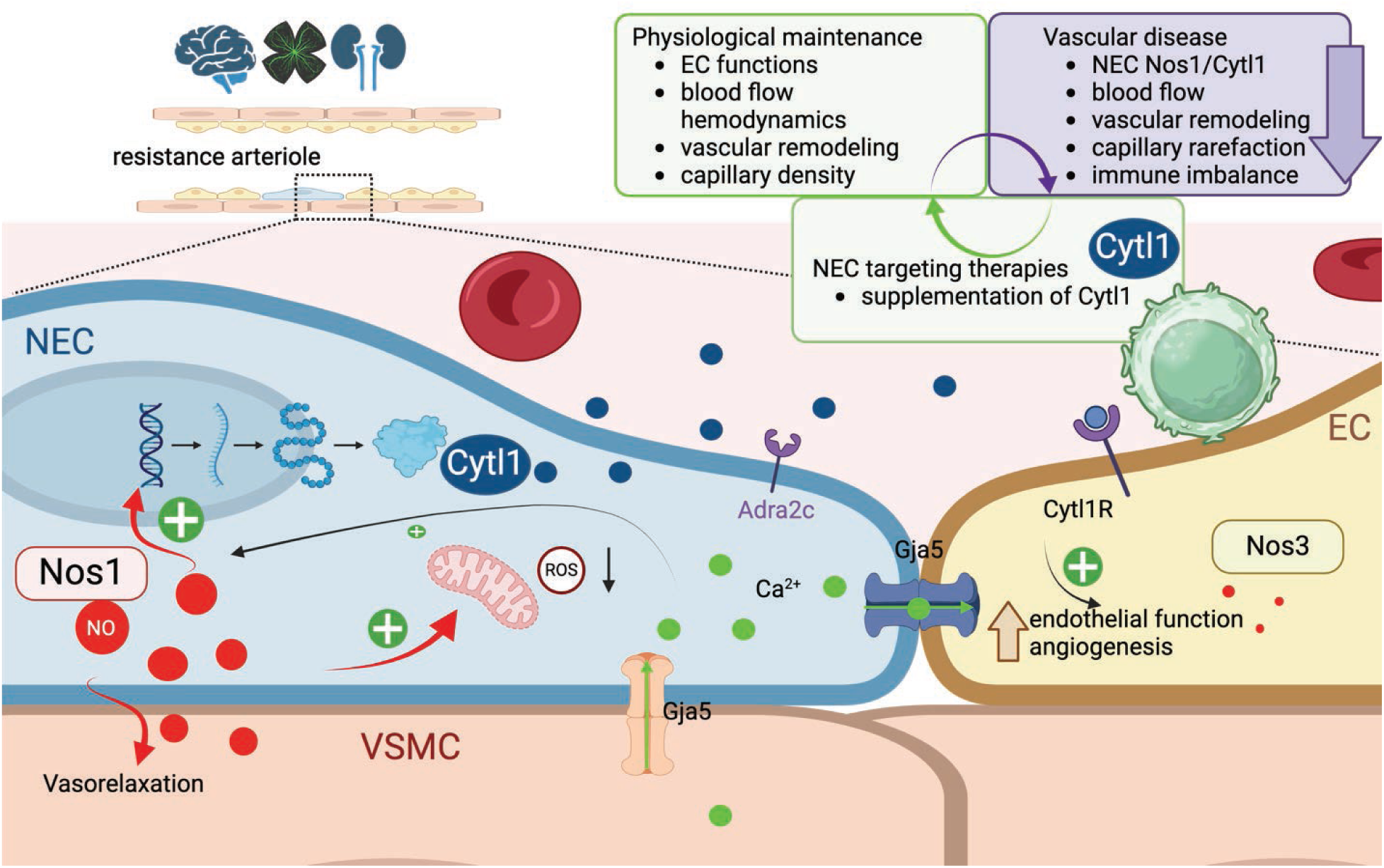
Schematic illustration of NEC functions.

## Discussion

This study discovered NECs, a permanent and neuronally differentiated endothelial cell subtype in small resistance arterioles of multiple organs and identified their key roles in the regulation of blood flow, vascular permeability and density, and immune cell homing in the local tissue microenvironment of organs in which they reside. The molecular signature of NECs including their top biomarkers Nos1 and Cytl1 suggested their biological functions in vasodilation, angiogenesis, mechanosensing, synaptic signaling, neurogenesis, and tissue inflammation, which were subsequently and functionally validated in vivo. The presently uncovered NEC-mediated endothelial mechanisms may represent a new paradigm in vascular biology and in the physiological maintenance of organ blood flow, tissue organization and homeostasis, and immune balance. NEC biomarkers were translated to the human condition, functionally linked to vascular and immune pathologies in the brain and kidney and may be targeted for therapeutic development. NECs were found with the highest density in the three organs (brain>retina>kidney, Fig. 1) that exhibit the highest metabolic and blood flow demand per organ weight^26^, the best blood flow autoregulation efficiency^27^, and are primarily affected by the development of microvascular complications in aging and diabetes^28^. Localization in small resistance arterioles in these three organs, and their vasoactivity and blood flow regulatory functions are consistent with NECs’ role in providing highly sensitive and efficient blood flow autoregulation. In addition, these anatomical and functional features suggest the importance of metabolic and hemodynamic cues in NEC development. In addition, NECs were found in the heart and pancreas, however with much lower density (Supplemental Fig. 1), suggesting NECs are also relevant to cardiovascular and endocrine physiology and body metabolism, and potentially to heart disease and diabetes.

The more elongated cell morphology of NECs compared to ECs may be explained by their exposure to higher shear stress at their predominant localization at branching segments of resistance arterioles^29^. A combination of structural, molecular, and functional features of NECs suggests their neuronal differentiation, including their long cell processes, numerous small membrane protrusions for synapse-like endo-endothelial or myoendothelial junctions (Fig. 1C), their molecular fingerprint with Nos1 and sensory and synaptic signaling profile (Figs. 2-3).

The present study developed and applied a comprehensive NEC research toolbox including transgenic mouse models with NEC reporter, NEC transcriptome, genetic silencing of top NEC factors in vivo, optogenetic tools in combination with intravital multiphoton microscopy (MPM), and injectable recombinant NEC-derived proteins. Optogenetic approaches provided proof-of-principle and confirmed the key regulatory role of NECs in the maintenance of organ blood flow (Fig. 4). The functional expression of Adra2c in NECs (Figs. 2D-F, 3C-E, Supplemental Fig. 4C) suggest the role of NEC adrenoceptors in stress-induced redistribution of regional blood flow. This is consistent with NEC-mediated direct vasodilator actions of norepinephrine to counterbalance vascular smooth muscle-mediated vasoconstrictions and maintain organ blood flow, similarly to the actions of angiotensin II on ECs^25^. Silencing of the top NEC factors Nos1 and Cytl1 resulted in a complex vascular phenotype resembling endothelial dysfunction, including severe vasoconstrictions, reduced blood flow and capillary density, increased vascular permeability and immune cell homing (Fig. 5). This phenotype further confirms and functionally validates the above key vascular NEC functions, and the role of NEC Nos1 and Cytl1 as novel major regulators of vascular and tissue homeostasis in the brain, retina, and kidney. Specifically, the present study identified Cytl1 as a novel and highly potent anti-inflammatory factor that contributes to the maintenance of immune balance in the local tissue environment. Cytl1 is a small 15kDa secreted protein which was initially discovered in CD34^+^ haematopoietic stem cells with proangiogenic properties, triggering the sprouting of mature endothelial cells largely independent of Vegfa^30^. Consistent with the presently identified anti-inflammatory function, Cytl1 was found to play immunomodulatory roles in cartilage homeostasis and osteoarthritis development^31^. Cytl1 may be used in future therapeutic development for a spectrum of neurological and kidney diseases that are based on vascular and immune pathologies.

Altogether, molecular and functional clues suggest that NECs are important and specialized sensory and regulatory ECs localized in resistance arterioles of select organs that are exposed to the highest mechanical and metabolic stress and require the most efficient blood flow autoregulation and vascular maintenance. NECs play key roles in peripheral tissue maintenance via their numerous inter-related functions that include the regulation of vasoactivity, blood flow, angiogenesis and capillary density, ECM and tissue remodeling, and inflammation.

## Methods

### Sex as a biological variable

Our study examined male and female animals, and similar findings are reported for both sexes.

## Animals

Male and female, 6-12 weeks old C57BL6/J or BalbC mice (Jackson Laboratory, Bar Harbor, ME) were used in all experiments. The presently used transgenic Cdh5(PAC)-CreERT2 mice^32^ were provided by academic investigator Ralf Adams (University of Münster, Münster, Germany, Cancer Research UK Scientist via Cancer Research Technology Limited). Transgenic mouse models with the expression of various fluorescent reporter proteins were generated by intercrossing Cre-ER^T^^2^ mice with flox mice as listed in the table below.

| Mouse Model | Cre-ER<sup>T2</sup> | Reporter | Induction |
| --- | --- | --- | --- |
| Cdh5-Confetti | *Cdh5*(PAC)<sup>32</sup> | *(Gt(ROSA)26Sortm1(CAG-Brainbow2.1)Cle/J)*<sup>33</sup> | tamoxifen via oral gavage once |
| Cdh5-GT | *Cdh5*(PAC)<sup>32</sup> | *Gt(ROSA)26Sortm4(ACTB-tdTomato,-EGFP)Luo*<sup>34</sup> | tamoxifen via oral gavage once |
| Nos1-GFP | *Nos1*<sup>35</sup> | *Gt(ROSA)26Sortm4(ACTB-tdTomato,-EGFP)Luo*<sup>36</sup> | prolonged tamoxifen |
| Nos1-GT | *Nos1*<sup>35</sup> | *B6;129S6-Polr2aTn(pb-CAG-GCaMP5g,-tdTomato) Tvrd/J*<sup>34</sup> | prolonged tamoxifen |
| Nos1-Ai27 | *Nos1*<sup>35</sup> | *RCL-hChR2(H134R)/tdT-D*<sup>37</sup> | prolonged tamoxifen |
| Nos1-Ai39 | *Nos1*<sup>35</sup> | *RCL-eNpHR3.0/EYFP*<sup>37</sup> | prolonged tamoxifen |

For conventional Cre/lox induction, tamoxifen was administered 75 mg/kg by oral gavage once. For prolonged tamoxifen induction, animals received ip injection of tamoxifen (75 mg/kg) for a total of four times once per week for 4 weeks, and tamoxifen containing diet (TD.130856, Inotiv Teklad, West Lafayette, IN) for 4 days on the first week of induction resulting in cell specific expression of reporters. A tamoxifen washout period of >2 weeks was always used. Some mice received acute iv. administration of norepinephrine (NE, 1mg/kg BW), selective Nos1 inhibitor 7-NI (20mg/kg ip., N778, Sigma Aldrich, St. Louis, MO), acute (20μg/kg BW iv.) or chronic (20μg/kg BW daily for 5 consecutive days sc.) administration of mouse recombinant Cytl1 injection.

### Adeno-Associated Virus Delivery of shRNA for genetic silencing

The endothelial targeting adeno-associated virus (AAV) construct, AAV2–QUADYF pAAV[shRNA]-CAG>EGFP-U6>mNos1 (Vector ID:VB240708-1385trz) or pAAV[shRNA]-EGFPU6>mCytl1 (Vector ID: VB900145-1533vjz) was used to knock down the expression of Nos1 or Cytl1 in NECs, respectively^38^. A scrambled shRNA control AAV2-QUADYF virus construct (Vector ID: VB010000-9489hhg) expressing GFP in target cells was used as the negative control. All constructs were obtained from VectorBuilder (Chicago, IL). Animals were randomized into control and knock down groups (n=8 mice in each group) 1 week before AAV administration. AAV constructs were suspended in sterile PBS and administered by retro-orbital injection in anesthetized animals at 8 weeks of age. AAV dose of 2×10^11^ genome copies was used as established earlier^38^. In vivo MPM imaging of the brain and kidneys was performed at the end of 2 weeks of follow-up period. Mice were euthanized after intravital imaging, and brain and kidney tissues were harvested.

### Unilateral ureteral obstruction

Unilateral ureteral obstruction (UUO) was performed as described before^39^. Briefly, between 6 and 8 weeks of age, the animals were anesthetized with isoflurane, and after a midline laparotomy the left ureter was exposed and ligated 3 times. Successful ligation was confirmed by the hydronephrotic distention of the kidney at the time of imaging (4 weeks after surgery). Control animals underwent sham operation. Eight mice were allocated to each of the 2 groups (control and UUO) via simple randomization.

### Intravital imaging using multiphoton microscopy (MPM)

Surgical implantation of a dorsal abdominal imaging window (AIW) above the left kidney or cranial imaging window (CIW) was performed on Nos1-Ai27, Ai39, GT, and Cdh5-GT mice using aseptic surgical procedures as described recently^25,40^. For MPM imaging, animals underwent brief anesthesia sessions using 1%–2% isoflurane and the SomnoSuite low-flow anesthesia system (Kent Scientific, Torrington, CT). For acute kidney imaging experiments under continuous anesthesia (Isoflurane 1-2% inhalant via nosecone), the left kidney was exteriorized through a flank incision. Mice were placed on the stage of the inverted microscope with the exposed kidney mounted in a coverslip-bottomed chamber bathed in normal saline and maintained as described previously^39,41,42^. Alexa Fluor 680-conjugated albumin (ThermoFisher, Waltham, MA) was administered iv. by retro-orbital injections to label the circulating plasma (30 µL iv. bolus from 10 µg/ml stock solution). The images were acquired using a Leica SP8 DIVE multiphoton confocal fluorescence imaging system with a Leica 25× water or 63× glycerine-immersion objective (numerical aperture (NA) 1.3) powered by a Chameleon Discovery laser at either 760 nm, 960 nm, 1100 nm (Coherent, Santa Clara, CA) and a DMI8 inverted microscope’s external Leica 4Tune spectral hybrid detectors (emission at 460-480nm for CFP, 510-530 nm for eGFP and GCaMP5, 550-570 nm for YFP, 580-600 nm for tdTomato, 600-620 nm for RFP) (Leica Microsystems, Heidelberg, Germany). The potential toxicity of laser excitation and fluorescence to the cells was minimized by using a low laser power and high scan speeds to keep total laser exposure as minimal as possible. Image acquisition (12-bit, 512×512 pixel) consisted of only one z stack or time series per tissue volume (<2-10 min), which resulted in no apparent cell injury. For tissue volume and cell density analysis fluorescence images were collected in volume series (xyz, 1 s per frame) with the Leica LAS X imaging software and using the same instrument settings (laser power, offset, gain of all detector channels). Maximal projections from *Z*-stacks were used to count and compare GFP^+^ or Confetti^+^ cell number in the same tissue volume. In the Confetti mouse models clonal or unicolor tracing units were defined as numerous directly adjacent individual cells that featured the same Confetti color combination. All ten possible Confetti color combinations were observed as described before^41^. The counting of GFP^+^ or Confetti^+^ cells and clones was facilitated by standardized image thresholding using ImageJ (NIH), Leica LAS X (Leica Microsystems Inc.), and cell-counting algorithms of Imaris 10.2 3D image visualization and analysis software (Oxford Instruments, Abingdon, UK) for intravital imaging *Z*-stacks. Clone frequency (the percentage of clones formed by 1, 2, 3-6, or >7 individual Confetti cells relative to all clones (100%)) was analyzed using mixed-effect analysis with two-way ANOVA^24^. Imaris 10.2 Surface and Filament tracing module was used to quantify capillary volume/tissue volume, and vessel segment diameter, respectively. To study dynamic changes in intracellular calcium signaling in ECs and NECs, fluorescence images were collected in time series (xyt, 526 ms per frame) with the Leica LAS X imaging software and using the same instrument settings (laser power, offset, gain of both detector channels) and analyzed as described before^24^. The strong cell-specific tdTomato fluorescence signal^42^ and high-resolution MPM imaging allowed for easy identification of single cell bodies and processes^43^. In the optogenetic mouse models, NEC specific stimulation (Nos1-Ai27) or inhibition (Nos1-Ai39) was achieved via exposing single NECs (identified by the genetic expression of tdTomato (Ai27) and EYFP (Ai39)) with blue light (470 nm, Ar laser) or red light (590 nm, He-Ne laser, same power), respectively, using ROI scanning and 1 minute sequences of 1 Hz square pulse trains for intermittent light exposure (Leica Microsystems Inc). Endogenous circulating immune cells were labeled by iv injection of 20 µl anti-CD45-Alexa Fluor 488 antibodies (30-F11, BioLegend, San Diego, CA). MPM imaging was always performed at the same time of day.

## Blood pressure measurements

Systolic blood pressure was measured by tail-cuff plethysmography (Visitech BP-2000, Visitech System Inc.) in trained animals as previously described^44^.

### Tissue clearing and light sheet imaging

Whole organ tissue clearing of various organs including the mouse brain, kidney, heart, lung, pancreas, liver, skeletal muscle, skin, and visceral fat was performed using the Shield protocol for tissue preservation^45^ and Clear+ tissue clearing technique according to manufacturer’s instructions (Passive Clearing Kit, LifeCanvas Technologies, Cambridge, MA). Light sheet imaging was performed using Miltenyi/LaVision Biotech Blaze Ultramicroscope with MI Plan 4x objective (numerical aperture 0.35), 2.4x magnification, detection with 4.2 Megapixel sCMOS camera, and laser lines 488 nm and 561 nm. Image processing and 3D reconstruction was performed using Imaris 10.2. (Oxford Instruments; Abingdon, UK).

### Tissue processing and immunofluorescence

Immunofluorescence detection of proteins was performed as described previously^44^. Briefly, cryosections were cut at 25 µm, washed with 1x PBS. Paraffin tissue blocks were sectioned to 8 µm thick. For antigen retrieval, heat-induced epitope retrieval with Sodium Citrate buffer (pH 6.0) or Tris-EDTA (pH 9.0) was applied. To reduce non-specific binding, sections were blocked with normal serum (1:20). Primary and secondary antibodies were applied sequentially overnight at 4° C and 2 hours at room temperature.

### RNAscope (mRNA Fluorescence In Situ Hybridization) and Immunofluorescence

Nos1 and Cytl1 mRNA detection combined with CD31 immunostaining in mouse brain and kidney sections were manually carried out using the RNAscope Multiplex Fluorescent v2 Assay with TSA Vivid Dyes (323 280; Advanced Cell Diagnostics, Newark, CA), and the RNA-Protein Co-Detection Ancillary Kit (323 180; Advanced Cell Diagnostics) as described recently^25^. Briefly, 4 μm formalin-fixed, paraffin-embedded slides were used. RNAscope Co-Detection Target Retrieval reagent was used for antigen-retrieval. Primary (CD31, 77 699; Cell Signaling Technology) and secondary antibodies (A11034, Invitrogen), and RNAscope™ Probe-Mm-Nos1 or Probe-Mm-Cytl1 (437651, 817241, respectively, Advanced Cell Diagnostics) and RNAscope Multiplex Fluorescent Detection Reagents were applied sequentially. Slides were counterstained with 4’,6-diamidino-2-phenylindole and mounted. All slides were imaged using a Leica TCS SP8 (Leica Microsystems) confocal/multiphoton laser scanning microscope system. The density of Nos1 and Cytl1 in single NECs identified by CD31 labeling was quantified using ImageJ (NIH) as described before^24^.

### Endothelial cell isolation

ECs and NECs with various labeling (genetically expressed tdTomato and GFP, Anti-CD31-Alexa700) were isolated from freshly harvested mouse brain and kidney. Tissues were minced into 1 mm^3^ pieces, digested using Hyaluronidase and Liberase enzyme combination (concentration, 2 mg/mL and 2.5 mg/mL, respectively, Sigma-Aldrich). The digested tissue was gently passed through a sequence of 100 µm, 70 µm, and 40 µm strainer three times, and washed with PBS. After digestion, ECs (tdTomato/Anti-CD31-Alexa700) and NECs (GFP) were isolated based on their genetic reporter expression or CD31 labeling by using BD SORP FACSYMPHONY S6 cell sorter equipped with 6 lasers (355, 405, 445, 488, 552, and 638 nm), and excitation wavelengths 488, 552 and 638 nm in sterile conditions. Data was analyzed using FlowJo (Tree Star, Ashland, OR).

### RNA sequencing and bioinformatics

Single cell RNA sequencing was prepared using 10x Genomics 3’ v3.1 (cat# 1000092) following manufacturer’s protocol as described before^24^. Samples were parsed into single cells using 10x Genomics Chromium Controller and libraries were simultaneously prepared. Prepared single cell RNA sequencing libraries were sequenced on the Illumina Novaseq6000 platform at a read length of 28×90 and read depth of 100,000 reads/cell for 100K cells.

Alignment and gene quantification of single-cell RNA sequencing data were performed using Cellranger v7.2 with default settings and the mouse mm10 2020-A dataset as reference. Cells exhibiting more than 5% of mitochondrial DNA content and fewer than 500 nFeature_RNA were excluded. Additionally, clusters primarily composed of low-quality cells were excluded. Gene counts were normalized and scaled to account for the total unique molecular identifier counts per barcode using Seurat v5.0.11. All subsequent processing steps followed standard guidelines in the Seurat package^46^, and as previously reported^47^. Cell clustering was achieved using the FindNeighbors function, followed by clustering with the FindClusters function. Dimensionality reduction was performed using Uniform Manifold Approximation and Projection (UMAP) through the RunUMAP function.

Classification of distinct cell types was guided by established cell-type markers from endothelial cell types in the mouse brain^1,5^ and kidney^1,48^. The identification of cell type–specific marker genes was performed using the FindMarkers function, with all parameters set to default. Genes with Bonferroni correction adjusted p-value <0.05 were considered marker genes. The cell types for the brain EC dataset were classified based on established markers^1,5^ for large artery, artery, artery shear stress, capillary arterial, capillary, capillary venous, large vein, choroid plexus. Cell types for kidney EC were classified based on established markers^1,48^ for arteriole, arteriole/efferent, capillary, glomeruli and venules. Brain and kidney EC datasets were subdivided into Nos1^+^ and Nos1^-^ arteries based on a gene expression threshold (brain: Nos1 > 5; kidney Nos1 > 4). FindMarkers function was used to identify markers between Nos1^+^ and Nos1^-^ arteries. Downstream analysis was performed including Gene Set Enrichment analysis (GSEA) using R package clusterProfiler^49^ using the default settings. Additionally, shared and differentially expressed genes in Nos1^+^ arteries in brain and kidney datasets were identified and over representation analysis (ORA) was performed with clusterProfiler6 using the default settings.

### Statistical methods

Data represent average ± SEM and were analyzed using Student’s t-tests (between two groups), or two-way (mixed-effect) or one-way ANOVA (for multiple groups) with post-hoc comparison by Dunnett’s, or Tukey’s, or Sidak’s tests as appropriate. P<0.05 was considered significant. Statistical analyses were performed using GraphPad Prism 9.0c (GraphPad Software, Inc.).

### Study approval

All animal protocols were approved by the Institutional Animal Care and Use Committee at the University of Southern California.

## Acknowledgments

This work was supported by a Carl W. Gottschalk Research Scholar Grant by the American Society of Nephrology and a Dean’s pilot funding award from the Keck School of Medicine at the University of Southern California (GG), and by US National Institutes of Health grants DK064324, DK123564, DK135290 and S10OD021833 (JPP).

## Author contributions

G.G. and J.P.P. designed the study, performed experiments, analyzed the imaging data, and wrote the manuscript. R.R. and Y.C. analyzed transcriptome data, S.W.R, J.A.J, and S.F. acquired and analyzed imaging data. A.B.C, A.I., G.T., S.D. performed experiments and made substantial contributions to acquire and analyze data. B.V.Z. provided guidance regarding brain imaging studies. All authors approved the final version of the manuscript.

