## Supplemental Material for "Neuronally differentiated endothelial cell subtype regulates organ blood flow and immune balance"

**This file contains the following Supplemental Materials:**

**Supplemental Figures 1-4**

**Supplemental Movies 1-3**

**Supplemental Table 1**

### Supplemental Figures

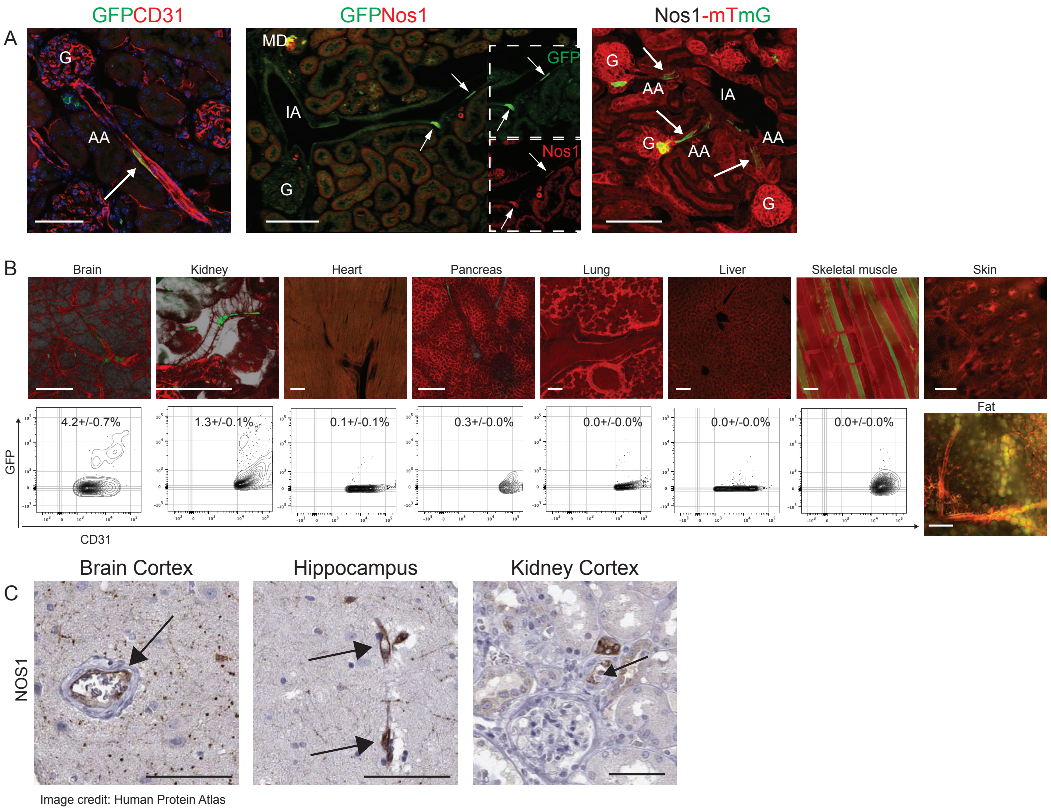


**Supplemental Figure 1. Additional features of NEC genetic labeling, localization, and organ distribution. (A)** Immunofluorescence double labeling of the genetic NEC marker GFP (green) with the endothelial marker CD31 (red) and their co-localization in NECs (yellow) confirming NECs as an endothelial subtype (left, arrow). Double labeling of GFP (green) and Nos1 (red) in kidney section of a 5 months old Nos1-mTmG mouse that received a single tamoxifen induction at 4 weeks of age (center). Insets show green and red channels separately. Note the co-localization of GFP and Nos1 in NECs (yellow, arrows) and in the macula densa (MD, positive control). Native fluorescence image from a frozen kidney section of a Nos1-mTmG mouse that received continuous tamoxifen induction for 8 weeks (right). Note the presence of a few scattered NECs in afferent arterioles (AA)(green, arrows). Cell nuclei are stained blue with DAPI. AA: afferent arteriole, G: glomerulus, IA: interlobular arteriole. **(B)** Representative native fluorescence images of fixed tissue sections from multiple organs of Nos1-mTmG mice (NECs are green, all other cells are red) with corresponding flow cytometry data and NEC density values underneath (n=4 each). **(C)** NOS1 immunohistochemistry in human kidney and brain tissue sections identify NECs in small arterioles (arrows, images from the Human Protein Atlas). Bars are 100 µm.

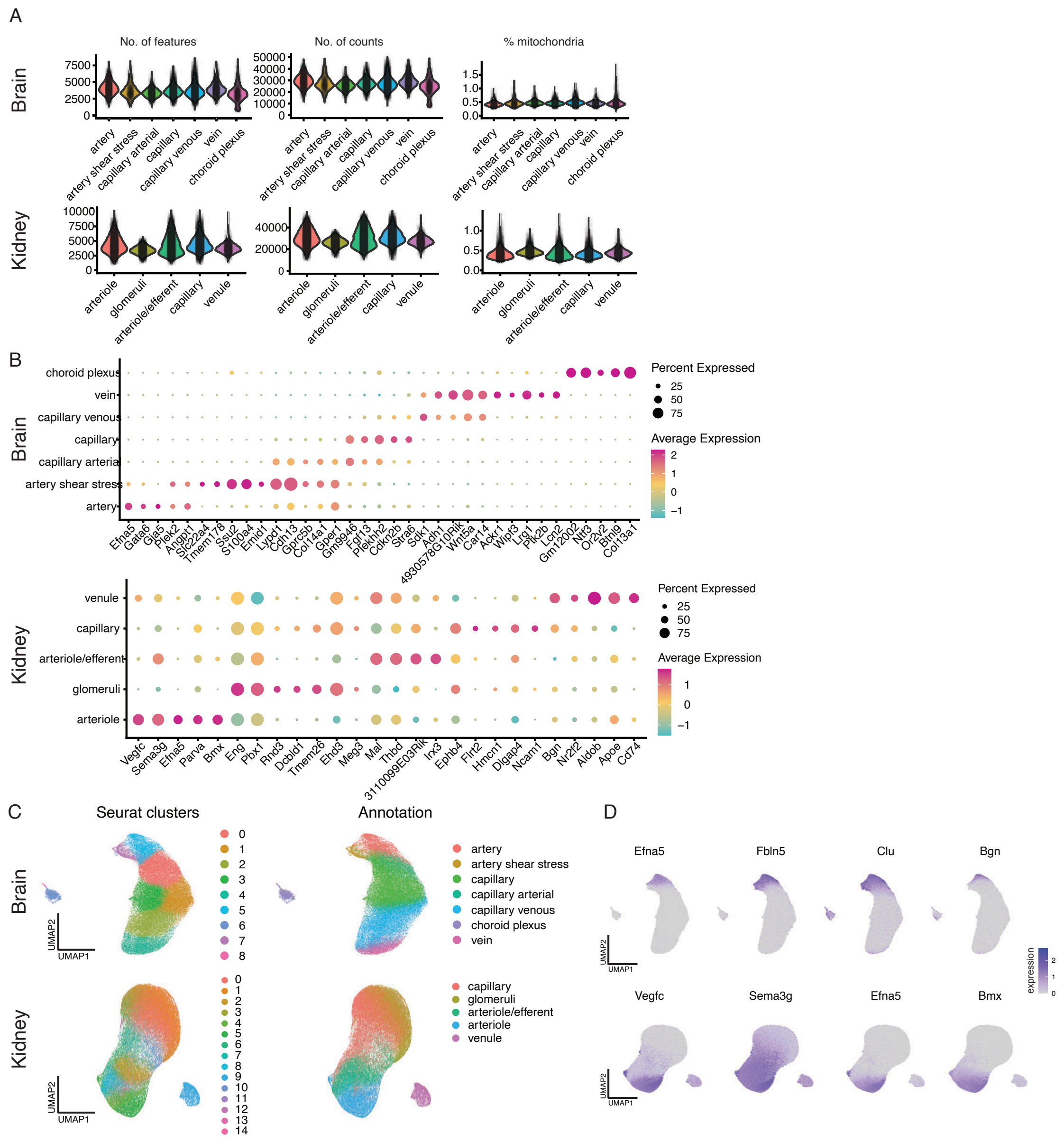


**Supplemental Figure 2. Quality controls and processing of scRNA sequencing and bioinformatics. (A)** Validation of the number of features, gene counts, and mitochondrial DNA content according to arterio-venous zonation. **(B)** Classification and marker identification of the main arterio-venous segment ECs based on the indicated established markers. **(C)** Clustering of ECs and annotation of arterio-venous zonation according to Seurat. **(D)** Expression of the applied representative arterial EC markers.

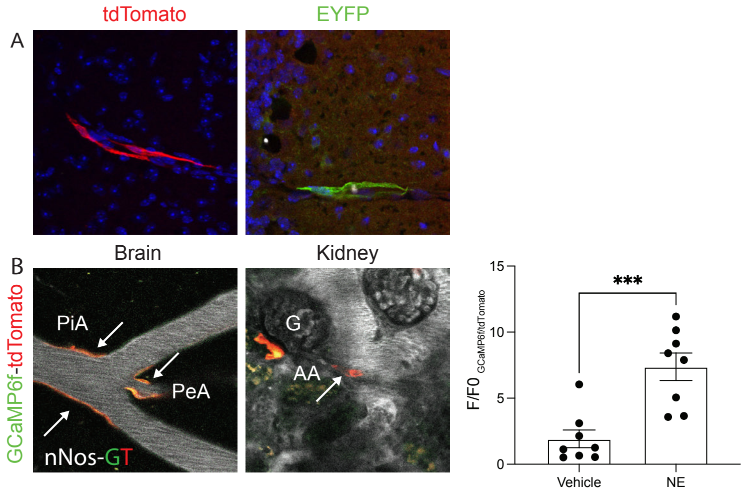


**Supplemental Figure 3. Model validation and intravital MPM imaging of NEC vasoactive functions. (A)** Validation of the NEC-Ai27 and NEC-Ai39 mouse models based on the detection of the membrane targeted Nos1-driven expression of the genetic reporters tdTomato (red, NEC-Ai27 model, left) or EYFP (green, NEC-Ai39 model, right) in frozen brain tissue sections. **(B)** Representative intravital images of NECs in mouse brain (left) and kidney (center) resistance arterioles in the NEC-GCaMP6f/tdTomato (GT) mouse model. Note that NECs (orange, arrows) express the calcium sensitive green fluorescent protein GCaMP6f and the red calcium insensitive fluorescent protein tdTomato. Statistical summary of changes in NEC calcium signaling (normalized to baseline) in response to acute iv administration of norepinephrine (NE, 1mg/kg BW) compared to vehicle (right) (n=8). PiA: pial artery, PeA: penetrating arteriole, AA: afferent arteriole, G: glomerulus. Data are mean ± SEM, ***$P<0.001$ with t-test.

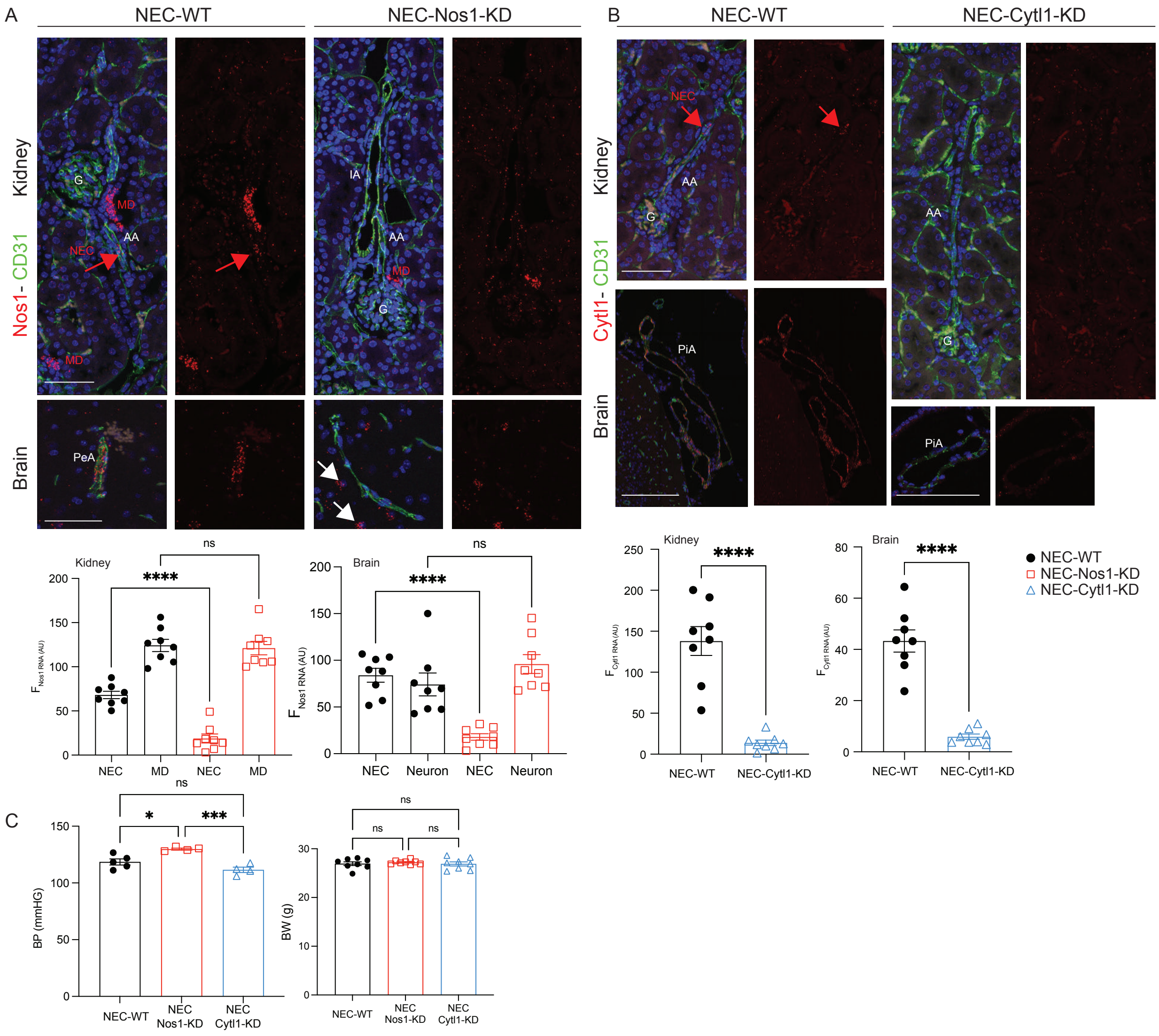


**Supplemental Figure 4. Validation and quantification of AAV2-QUADYF-mediated NEC Nos1 and Cytl1 gene silencing in vivo. (A-B)** Representative fluorescence images of double-labeling endothelial CD31 (green, immunofluorescence) with either Nos1 (red, RNAscope)(A) or Cytl1 (red, RNSscope) RNA (B) in pial (PiA) and penetrating (PeA), and afferent arterioles (AA) and interlobular arterioles (IA) in fixed brain and kidney tissue sections from AAV2-QUADYF-control RNA or AAV2-QUADYF-shRNA treated mice. Quantification of Nos1 and Cytl1 gene expression in control RNA and shRNA-treated mice (n=4). Nos1$^{+}$ macula densa (MD) or neurons served as positive controls. **(C)** Statistical summary of systolic blood pressure (BP, left) and body weight (BW, right). n=4-8. ns: not significant, *P<0.05,***P<0.001, ****P<0.0001with t-test or ANOVA with Tukey's or Sidak's test.

**Supplemental Movie 1. Localization of NECs in the mouse brain vasculature.** Representative 3D volume reconstruction of the motocortex region of a fixed, optically cleared whole-mount Nos1-mTmG mouse brain tissue based on light-sheet fluorescence imaging. Note the presence of a few scattered NECs (green) in pial and penetrating arterioles in the brain, while all other cell types are labeled by the genetic reporter tdTomato (red).

**Supplemental Movie 2. Intravital MPM imaging of the hemodynamic effects of optogenetic NEC stimulation in a mouse brain.** Representative time-lapse intravital MPM images of the vasodilator effect of blue light stimulation of a single NEC (identified by the genetic Nos1-driven tdTomato reporter, red, arrow on the left) using ROI scanning in a NEC-Ai27 mouse brain resistance arteriole. Plasma was labeled by iv injected albumin Alexa Fluor 680 (greyscale). Note the substantial vasodilatation of the entire penetrating arteriole upon blue light stimulation.

**Supplemental Movie 3. Intravital MPM imaging of the effects of acute Cytl1 administration in the mouse brain.** Representative time-lapse intravital MPM images of the hemodynamic changes of pial arteriole branches in the brain of a Cdh5-GCaMP6f/tdTomato (GT) mouse in response to iv injected mouse recombinant Cytl1 (20 μg/kg BW). Plasma was labeled by iv injected albumin Alexa Fluor 680 (greyscale). Note the substantial vasodilatation of the entire pial arteriole network upon Cytl1 administration. NEC/EC calcium signals are visible based on the expression of the calcium sensitive green fluorescent protein GCaMP6f and the red calcium insensitive fluorescent protein tdTomato.

**Supplemental Table 1. List of the top differentially expressed genes (DEGs) in NECs in the mouse brain.** The top 200 DEGs in brain NECs and the top 40 DEGs that are shared between the brain and kidney are listed with their statistical summary.
